# Mycophenolate mofetil increases susceptibility to opportunistic fungal infection independent of lymphocytes

**DOI:** 10.1101/131540

**Authors:** Rory H. Gibson, Robert J Evans, Richard Hotham, Aleksandra Bojarczuk, Amy Lewis, Ewa Bielska, Robin C. May, Philip M. Elks, Stephen A. Renshaw, Simon A. Johnston

## Abstract

Anti-proliferative agents that target lymphoid cells are common immunosuppressive agents used in the treatment of diverse autoimmune, graft versus host and inflammatory diseases. Mycophenolate mofetil (MMF) is an anti-proliferative agent that targets lymphoid dependence on inosine monophosphate dehydrogenase for the *de novo* purine synthesis of deoxyguanosine triphosphate (dGTP) for DNA replication. Here we show that MMF has a distinct and specific *in vivo* effect on macrophages, in the absence of lymphoid cells. This results in increased macrophage cell death that is dependent on the depletion of cellular GTP, independent of DNA synthesis. Furthermore, the macrophage specific effect of MMF treatment causes an increase in susceptibility to the opportunistic fungal infection *Cryptococcus neoformans* by reducing phagocytosis and increasing the release of intracellular pathogens via macrophage lysis. Our study demonstrates the need for a better mechanistic understanding of immunosuppressive treatments used in clinical practice and of the specific infection risks associated with certain treatment regimens.

## Introduction

Mycophenolate mofetil (MMF) is an anti-proliferative agent used in immunosuppressive therapy. MMF was developed as a morpholinoethyl ester prodrug of mycophenolic acid to improve bioavailability upon oral administration (Lee et al., 1990). Clinical trials in the 1990s identified MMF as a new immunosuppressive treatment which resulted in its introduction as part of combination therapy with a calcineurin inhibitor and steroids (Sollinger et al., 1992), and MMF is now often used to avoid toxicity associated with other immunosuppressants (Hueso et al., 1998). MMF plays an integral role in solid organ transplant regimes, alongside its increasingly common use for autoimmune disease, and it is now the mainstay of treatment for diseases such as systemic lupus erythematosus (Appel et al., 2005; Karim et al., 2002). However, like all immunosuppressive regimens, MMF is associated with increased susceptibility to infection.

MMF is a potent, selective, non-competitive inhibitor of the inosine monophosphate dehydrogenase enzyme (IMPDH) enzyme. IMPDH plays a major role in the *de novo* purine synthesis pathway that generates guanosine triphosphate (GTP), and deoxyguanosine triphosphate (dGTP) for DNA synthesis. While lymphocytes are largely dependent on *de novo* synthesis, other cells can use a salvage pathway, forming purine monophosphates from free purine bases. Thus, MMF acts as an anti-proliferative agent with good specificity for lymphocytes.

Cryptococcus neoformans is an opportunistic fungal pathogen of humans that causes cryptococcal meningitis (CM) in the severely immunocompromised, notably those with HIV-coinfection, with CD4 counts <200cells/mm^3^ (Gibson and Johnston, 2015). CM is difficult to treat, with a high mortality, due to late presentation and the limitations of current anti-fungal agents, especially in resource limited settings. Therefore, it is vital that cryptococcal disease is identified and treated as early as possible (Boulware et al., 2014). While the overwhelming disease burden of cryptococcal infection is associated with HIV immunosuppression, there are a number of other immunosuppressed groups that have increased incidence of cryptococcosis (Kuo et al., 2016; Neofytos et al., 2010). Recently, study of immune phenotype of non-HIV associated CM has demonstrated clear differences in macrophage activation and activity in the absence of CD4 T cell differences (Panackal et al., 2015).

Following solid organ transplantation, MMF is used, often in preference to other anti-proliferative agents such as azathioprine (AZA), due to its higher potency. A study of MMF versus AZA following renal transplant demonstrated that there was more than a three-fold increase in the risk of opportunistic infection with MMF treatment (Meier-Kriesche et al., 1999). Furthermore, there was clear increase in fungal infection with MMF treatment over AZA and this was the only independent risk factor for fungal infection (Meier-Kriesche et al., 1999). A systematic review of MMF versus AZA concluded that associating pathogens with different immunosuppression regimens is severely hampered by a lack of consistent reporting, sometimes only reported at the level of viral, bacterial or fungal cause (Wagner et al., 2015).

There are specific case reports that identify cryptococcal infection in patients treated with mycophenolate (Kluger et al., 2009; Marques et al., 2013), and the Transplant-Associated Infection Surveillance Network (a consortium of 23 US transplant centres) identified a 12-month cumulative incidence of invasive fungal infection (IFI) as 3.1%, with cryptococcosis the cause of almost 10% of IFIs (Pappas et al., 2010). Furthermore, there is little or no available data on specific immunosuppressive regimens and anti-fungal prophylaxis in such patient groups, making it impossible to identify the mechanistic relationship between observed infections and patient susceptibility.

Innate immune cell function is critical in handling opportunistic infections and macrophages are essential for control of cryptococcal infection (Bojarczuk et al., 2016; Osterholzer et al., 2009). Both steroids and MMF are thought to act in large part through modulation of lymphocyte function but it has been proposed that some of their clinical effect might be through modulation of innate immune cell function (Brack et al., 1997; Jiang et al., 2012; Meagher et al., 1996; Shah et al., 2010; von Vietinghoff et al., 2011). Therefore, we hypothesized that MMF might directly impair innate immune cell functions, revealing new opportunities for addressing the immune defect in MMF treated patients. To address this hypothesis, we tested MMF in a model system where innate immunity operates without functional adaptive immunity, our larval zebrafish model of cryptococcosis. We could identify a clear defect in resistance to cryptococcal infection with MMF but not AZA treatment. Increased susceptibility was due to increased macrophage cell death, resulting in less phagocytosis early in infection and enhanced release of intracellular cryptococci following lysis of infected macrophages. Therefore, MMF has a direct effect on macrophage control of the opportunistic fungal pathogen *C. neoformans*, independent of any lymphoid mechanism, a finding that has significant implications for the risks of infection with MMF treatment and the role of lymphoid cell dysfunction in MMF immunosuppression.

## Materials and Methods

### Ethics statement

Animal work was carried out per guidelines and legislation set out in UK law in the Animals (Scientific Procedures) Act 1986, under Project License PPL 40/3574. Ethical approval was granted by the University of Sheffield Local Ethical Review Panel.

### Fish husbandry

*Nacre* (White et al., 2008) was used as our wild type strain. Two macrophage (*Tg(mpeg1:Gal4.VP-16)sh256;Tg(UAS:Kaede)s1999t* (Prajsnar et al., 2013) and *Tg(mpeg1:mCherryCAAX)sh378* (Bojarczuk et al., 2016) and one neutrophil (*Tg(mpx:GFP)i114*) (Renshaw et al., 2006) fluorescent transgenic zebrafish lines were used. Annexin V labelling was performed using the *Tg(b-actin2:Gal4-VP16)sh330;Tg-(UAS:secAnxV.YFP)sh308* line crossed with *Tg(mpeg1:mCherryCAAX)sh378* (van Ham et al., 2010). TNFα expression was measured using the TNF promoter BAC driving GFP line, *TgBAC(tnfa:GFP)pd1028*. Zebrafish strains were maintained per standard protocols (Dahm and Nusslein-Volhard, 2002). Adult fish were maintained on a 14:10-hour light/dark cycle at 28°C in UK Home Office approved facilities in the Bateson Centre aquaria at the University of Sheffield.

### *Cryptococcus neoformans* culture

The *C. neoformans* variety grubii strain H99 and its GFP-expressing derivative H99GFP have been previously described in zebrafish (Bojarczuk et al., 2016). In addition, we generated and characterised KN99α GFP-expressing and mCherry-expressing derivatives (Fig. S3; see below for details of construction). 2ml YPD (reagents are from Sigma-Aldrich, Poole, UK unless otherwise stated) cultures were inoculated from YPD agar plates and grown for 18 hours at 28°C, rotating horizontally at 20rpm. Cells were pelleted at 3300g, washed twice with PBS (Oxoid,Basingstoke, UK) and resuspended in 2ml PBS. Washed cells were counted with a hemocytometer and used as described below.

### Plasmids and Strains

*C. neoformans* serotype A, strain KN99 α was biolistically transformed (Voelz et al., 2010) with a plasmid pAG32_GFP (Voelz et al., 2010) encoding for green fluorescent protein (GFP) or with a plasmid pRS426H-CnmCherry encoding for a *C. neoformans* - codon optimized red fluorescent protein mCherry. Stable transformants were tested for their sensitivity against several stress conditions mimicking hostile environment inside the phagocytes (Fig. S4). Three independent experiments were carried out where the transformants grew up at 37°C and 5% CO_2_ without shaking in the presence/absence of H_2_O_2_ or NaCl mimicking oxidative or cell wall stresses, respectively (Voelz et al., 2010); Fig S4). After 24 hours of growth serial dilutions of surviving cells were plated onto YPD plates and colony-forming units (CFU) were counted after 0 and 48 hours growth at 25°C incubator. CFUs relative to time point 0 were calculated. (Fig. S4). The plasmid pRS426H-CnmCherry which contains the hygromycin resistance cassette and *C. neoformans* codon optimised mCherry gene under the *histone 3* promoter was generated using *in vivo* recombination in *S. cerevisiae* strain FY834 (*MAT*α *his3* Δ*200 ura3-52 leu2*Δ*1 lys2*Δ*202 trp1*Δ*63;*(Winston et al., 1995)) following published protocols (Knop et al., 1999; Raymond et al., 1999). Firstly, a shuttle yeast-*E.coli*-*C.neoformans* KN99α plasmid pRS426H was made.

### Primers

EB90) GAAGTCTATGGGCGCTAGCGTCAAATGCGATGCGTGGGC

EB10) GAAAAGTTGCGGCGCC AAGCTT TGTGATAAATGTACTGAGGTGG

EB11) CATTTATCACAAAGCTTGGCGCCGCAACTTTTCTTAGCTTGCTTTTTG

EB12) GATGTATGTGTTTAAACTATACGCGATTACAAGTATTTGTAG

EB13) GTAATCGCGTATAGTTTAAACACATACATCCCCTATACCGCATC

EB14) CTTTTTACCCAT ACGCGT GTTGGGCGAGTTTACTAATGG

EB15) GTAAACTCGCCCAAC ACGCGT ATGGGTAAAAAGCCTGAACTCAC

EB16) CCGCCTTCAC GCGGCCGC TTATTCCTTTGCCCTCGGACG

EB17) GGCAAAGGAATAA GCGGCCGC GTGAAGGCGGTAAGGGGTTAAT

EB24) CAGCAACAACCACCTCAGTACATTTATCACAAAGCTTATGGTCTCCAAGGGTGAGGAG

EB25) GTCTACAAAAAGCAAGCTAAGAAAAGTTGCGGCGCCTTACTTGTAGAGCTCGTCCATAC

Plasmid pRS426H was made using a backbone containing *ampicillin* resistance gene, *E.coli ori* and *URA3* selectable marker from a commercially available plasmid pRS426 digested with *KpnI* and *SacI*. In addition, pRS426H contains hygromycin resistance gene under a constitutive *actin* (CNAG_00483) promoter and a *trp1* (CNAG_04501; anthranilate synthase/indole-3-glycerol phosphate synthase/phosphoribosylanthranilate isomerase) terminator and a *histone 3* (CNAG_06745) promoter and terminator with a *HindIII* and *NarI* restriction sites between them as a potential site for introduction of a gene of interest. A 531 base pairs (bp) region of *histone 3* promoter was amplified from *C. neoformans* KN99α genomic DNA using primers EB9 and EB10. EB10 primer introduced two restriction sites: *HindIII* followed by *KasI* at the end of the promoter region allowing cloning a gene of interest at the end of the promoter. A 399 bp region of *histone 3* terminator was amplified from *C. neoformans* KN99α genomic DNA using primers EB11 and EB12. EB11 primer introduced two restriction sites: *HindIII* followed by *KasI* in a front of the terminator region allowing a potential site for introduction of a gene of interest in a front of the terminator. A 562 bp region of *actin* promoter was amplified from *C. neoformans* KN99α genomic DNA using primers EB13 and EB14. EB14 primer introduced *MluI* restriction site at the end of the promoter region allowing a potential change of a resistance gene. A 1029 bp region of *hygromycin* gene was amplified from pAG32_GFP (Voelz et al., 2010) using primers EB15 and EB16. The primers introduced two restriction sites: *MluI* in a front of the resistance gene (by EB15) and *NotI* at the end of the resistance gene (by EB16) allowing a potential change of the resistance gene. A 248 bp region of *trp1* terminator region was amplified from *C. neoformans* KN99α genomic DNA using primers EB17 and EB18. EB17 primer introduced *NotI* in a front of the terminator region allowing a potential replacement a resistance gene. Finally, the plasmid pRS426H-CnmCherry was obtained by introducing a gene encoding *C. neoformans* codon optimized mCherry amplified from pKUTAP (a kind gift from Peter Williamson, Laboratory of Clinical Infectious Diseases, NIAID, NIH, Bethesda, Maryland, United States of America) in between *histone 3* promoter and terminator of the plasmid pRS426H initially linearized with *HindIII* and *KasI* using primers EB24 and EB25.

### Drug treatments

Following infections at 2dpf, larvae were allocated into random treatment/control groups and placed into a 96 well plate. All treatments were by immersion. The following treatment/control groups were investigated (as described in the figures, legends and text): 0.5µM MMF, 0.05% dimethyl sulfoxide (DMSO), 1mM 1400W in E3, 200µM LNIL in E3 (Cayman Chemical) 5.7µM nor-NOHA in 0.1% DMSO (Cayman Chemical), 200µM Azathioprine in 0.16mM NaOH and 750µM 6-Thiogunanine in 0.9mM NaOH.

Mycobacterial infection and bacterial burden.

Infection experiments were performed using *M. marinum* strain M (ATCC #BAA-535), containing the pSMT3-Crimson vector. 150 colony-forming units (CFU) of bacteria were injected into the caudal vein at 2 dpf and bacterial burden measured at 3dpi as previously described (Elks et al., 2013).

### Fungal burden measurement

Zebrafish were infected with *C. neoformans* as we have described previously (Bojarczuk et al., 2016) with inoculum noted in figure legends. All injected zebrafish were incubated at 28°C until 3 days post infection. For imaging zebrafish were anaethetised with E3 containing 0.168 mg/mL tricaine. To assess fungal burden larvae were imaged using Nikon Ti-E with a CFI Plan Achromat UW 2X N.A. 0.06 objective lens, using Intensilight fluorescent illumination with ET/sputtered series fluorescent filter 49002 (GFP) (Chroma, Bellow Falls, VT, USA). Images were captured with Neo sCMOS, (Andor, Belfast, UK) and NIS-Elements (Nikon, Richmond, UK). The images produced show the extent of infection within the larvae, ranging from complete clearance to extensive fungal burden and dissemination. Images were exported as tif files and further analysis performed in ImageJ (Schneider et al., 2012). Images were individually cropped to remove the side of the 96-well or any bright debris or noise within the well. Pixels above the intensity corresponding to *C. neoformans* strain H99GFP were selected using a threshold. The same threshold was used for all images. Threshold images were converted to binary images and the number of pixels counted using the ‘analyse pixel’ function.

### Flow cytometery

To investigate the effect of MMF and Azathioprine on macrophage numbers we developed a flow cytometry method for dissociated larval zebrafish. Macrophage numbers were counted at 1, 2, and 3 days post treatment with MMF or AZA. Six larvae were dissociated over 1 hour in 500µL of trypsin with pipetting every 15 minutes to aid dissociation of larvae. Dissociated larvae were filtered through flow cytometry flow strainers with 700µL of PBS. We analysed macrophage *Tg(mpeg1:mCherryCAAX)sh378* and neutrophil (*Tg(mpx:GFP)i114*) numbers following treatment with MMF and AZA and their relevant vehicle controls. Flow cytometry was performed on the Atunne NXT (Applied Biosystems). Nacre wildtype strain was used to set negative population and macrophage and neutrophil populations counted from fluorescent population. Flow cytometry data were analysed with FlowJo (LLC, Oregon, USA).

### Time lapse imaging

The zebrafish line *Tg(mpeg1:Gal4.VP-16)sh256;Tg(UAS:Kaede)s1999t*, was infected with the cryptococcal strain KN99-mCherry. Following injection at 2dpf, zebrafish were randomly allocated into a MMF treatment group, and DMSO control, in a 96 well plate. At 1 day post infection zebrafish were anaesthetised by immersion in 0.168 mg/mL tricaine in E3 and immobilisation using in 0.5% low melting point agarose (containing either MMF, DMSO). Time lapse images were captured using a Nikon Ti-E with a CFI Plan Apochromat λ 10X, N.A.0.45 objective lens, a custom built 500 μm Piezo Z-stage (Mad City Labs, Madison, WI, USA) and using Intensilight fluorescent illumination with ET/sputtered series fluorescent filters 49002 (GFP) and 49008 (mCherry) (Chroma, Bellow Falls, VT, USA). Images were captured with Neo sCMOS, 2560□× □2160 Format, 16.6□mm x 14.0□mm Sensor Size, 6.5□μm pixel size camera (Andor, Belfast, UK) and NIS-Elements (Nikon, Richmond, UK). Time-lapses lasted 12 hours with 21 z-stack images 5μm apart, acquired every 4 minutes.

### TNFα reporter assay

Using the zebrafish transgenic *TgBAC(tnfa:GFP)pd1028* (Nguyen-Chi et al., 2015a) we measured the fluorescence signal from KN99-mCherry infected and uninfected larvae with and without MMF treatment. Larvae were imaged as described for fungal burden but for quantification each fluorescent area was segmented and its fluorescence intensity measured. Each of these measurements was then summed for each infection.

### Annexin V reporter assay

Annexin V positive macrophages assay was performed using the *Tg(b-actin2:Gal4-VP16)sh330;Tg-(UAS:secAnxV.YFP)sh308* line crossed with *Tg(mpeg1:mCherryCAAX)sh378*. Images were captured as described above for time lapse microscopy.

### High content imaging of host pathogen interactions

Measurement of macrophage and cryptococcal intracellular and extracellular numbers was completed as described previously in detail (Bojarczuk et al., 2016).

### Statistical analysis

Statistical analysis was performed as described in the results and figure legends. We used Graph Pad Prism 7 (v7.0a) for all statistical tests and plots.

## Results

### Mycophenolate mofetil increases susceptibility to cryptococcal infection in the absence of adaptive immunity

To test the hypothesis that mycophenolate mofetil (MMF) increases susceptibility to cryptococcal infection in the absence of adaptive immunity, we used our zebrafish larval model of cryptococcosis (Bojarczuk et al., 2016). We found that MMF increased cryptococcal fungal burden at 48 and 72 hours post infection (hpi; Fig. 1A-C) but had no effect on burden in the well-characterised bacterial zebrafish model of *Mycobacterium marinum* (Fig. S1). MMF is a member of a class of anti-proliferative drug that inhibits DNA synthesis through interfering with synthesis of GTP (Fig. 1F). AZA and 6-TG are two compounds that inhibit DNA synthesis by stalling nucleoside addition with 6-thioguanidine (Fig. 1F). In contrast, MMF results in a cellular depletion of GTP by directly blocking the synthesis of GMP (Fig. 1F; (Jagodzinski et al., 2004)). We found that azathioprine (AZA) and 6-thioguanine (6-TG) did not increase fungal burden up to 72hpi (Fig. D, E) and we concluded that the increased susceptibility to cryptococcal infection seen with MMF, and not with AZA and 6-TG, was due to the cellular depletion of GTP instead of an inhibition of DNA synthesis directly. We next examined the inflammatory activation of macrophages with MMF treatment.

**Fig 1.**
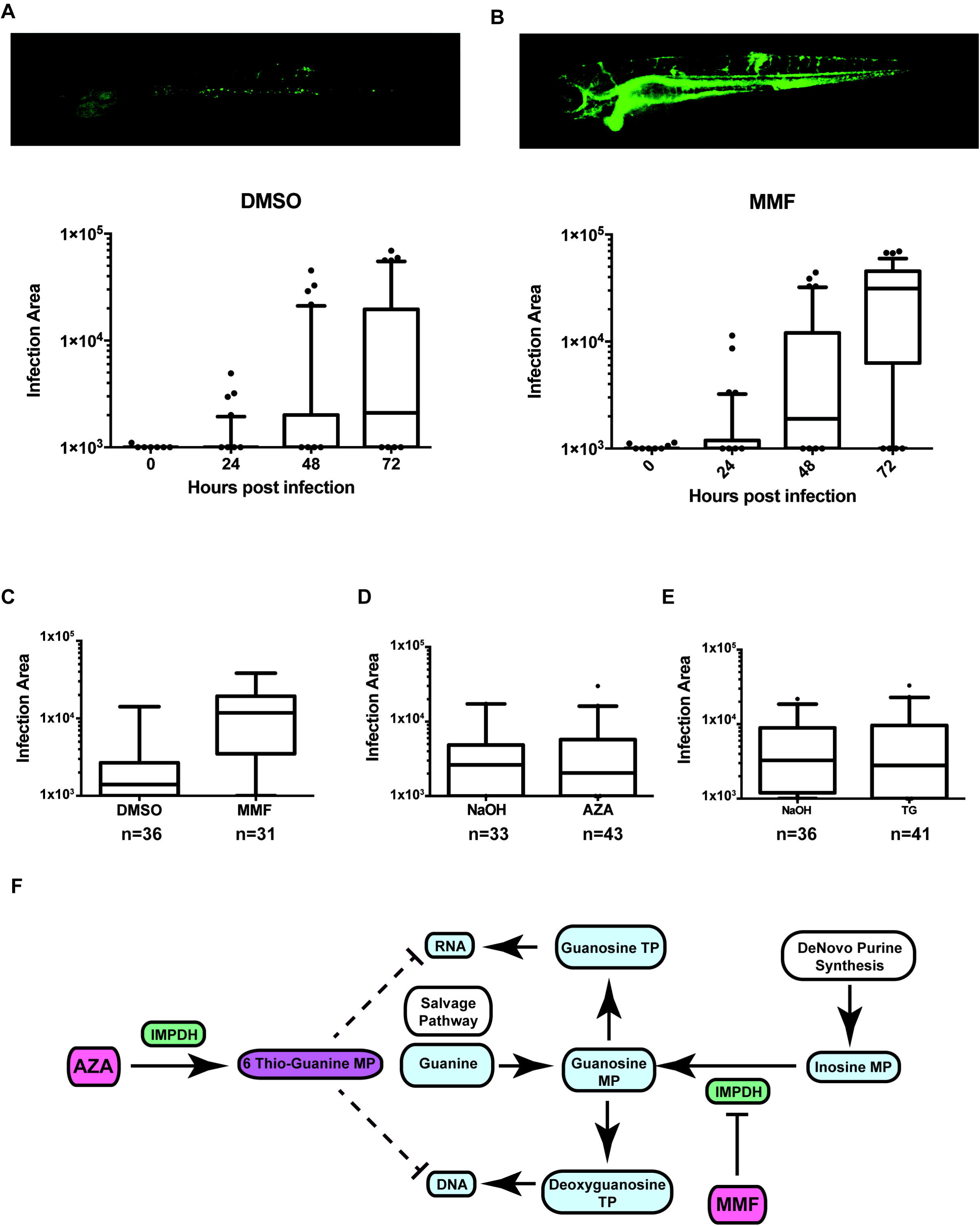
Treatment with Mycophenolate mofetil increased susceptibility to cryptococcal infection in the absence of adaptive immunity. A, B. Representative fluorescence image of the median H99GFP intensity for DMSO versus MMF. Box plots show increase in fungal burden over time. Whiskers are 5 and 95 percentiles with outliers. 48 hours post infection (hpi): median, DMSO= 568.5 MMF=1895, P=0.0005 and 72 hpi: median, DMSO=2102, MMF=29994, P<0.0001. 90 infections of 500cfu per group (30 per 3 biological repeats). **C.** Fungal burden at 72 hpi with 500cfu KN99αGFP infection. Median, DMSO=1485, MMF=8887, P<0.0001. Experiments with *C. neoformans* strain H99GFP and KN99alphaGFP gave indistinguishable results. **D, E.** Azathioprine and 6-Thioguanine do not affect susceptibility to cryptococcal infection. Drugs and concentration used 0.5µM MMF, 0.05% DMSO, 200µM Azathioprine in 0.16mM NaOH and 750µM 6-Thiogunanine in 0.9mM NaOH. **C-E.** n, is number of infections per 3 biological repeats. **F.** Simplified biochemical pathway of AZA and MMF mechanism. IMPDH, inosine monophosphate dehydrogenase.

### MMF inhibition of the iNOS pathway or TNFα **is not responsible for increased susceptibility to cryptococcal infection.**

Activation of macrophage inducible nitric oxide synthase (iNOS) by pro-inflammatory cytokines (principally interferon gamma (IFNg) and tumour necrosis factor alpha (TNFα) has been linked to the control and clearance of cryptococcal infection (Leopold Wager et al., 2014). Tetrahydrobiopterine (BH4) is an essential cofactor of iNOS and levels of BH4 depend on cellular GTP (Ionova et al., 2008). While IFNg is almost exclusively generated by lymphoid derived cells absent in our innate immune model, TNFα is produced abundantly by macrophages, is required for the protection against cryptococcal infection and is well characterised in zebrafish models (Nguyen-Chi et al., 2015b; Rayhane et al., 1999; Xu et al., 2016). Therefore, we tested the requirements for the iNOS pathway and production of TNFα in macrophage control of cryptococcal infection.

Using a fluorescent reporter of TNFα expression, macrophages generate significant levels of TNFα in response to *C. neoformans* as early as 24 hours post infection (Fig. 2A-E). TNFα was not expressed in macrophages at time of infection in infected larvae, and in uninfected groups at all time points (Fig. 2A-C). MMF treatment did not alter TNFα with or without infection (Fig. 2A-D). While there appeared to be an increase in TNFα expression between 24 and 72hpi in infected groups this was due to fewer, brighter areas at 72 hours and the total quantified fluorescence was the same at 24 and 72hpi (Fig. 2B-D). Using two inhibitors of iNOS (Elks et al., 2013) we found a modest but significant effect on fungal burden (Fig. 2F,G; 1400w, P=0.046; LNIL, P=0.014). The inhibitor 1400w displaces the BH4 cofactor from iNOS, and showed a lesser effect than the active site inhibitor LNIL (fold-change in median 1400w=1.5 vs LNIL=2.1). The small change observed with both inhibitors suggested that there was not a major role for the iNOS pathway in macrophage innate immune control of cryptococcal infection, and since these inhibitors would have direct effect on the iNOS pathway, in contrast to MMF, this suggested that this was not the mechanism of MMF action. Furthermore, we stimulated iNOS via inhibition of the enzymatic and signalling competitor arginase, using the drug NorNOHA. Arginase inhibition had no effect on fungal burden further supporting our conclusion that the iNOS pathway was not significantly active in macrophages during cryptococcal infection in the absence T-helper cell stimulation (Fig. 2H). In all cases, there was no interaction with MMF treatment (Fig. 2B-C; 1400w, P=0.98; LNIL, P=0.42; NorNOHA, P=0.33). Finding that there did not appear to be an inflammatory defect in macrophages treated with MMF we investigated changes in innate immune cell numbers.

**Fig 2.**
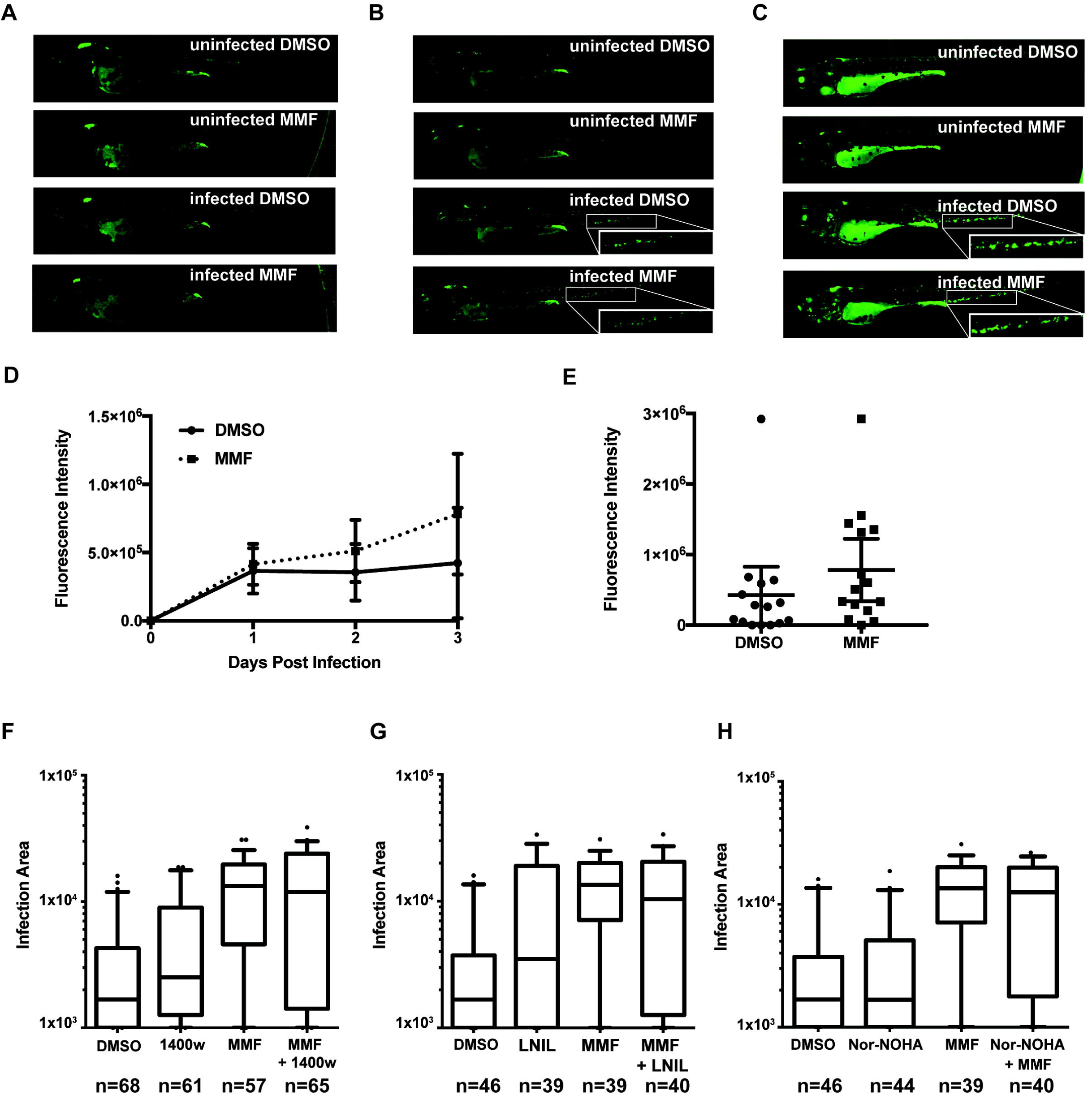
TNFα and iNOS responses of innate immune system were not modulated by MMF. **A.** TNF reporter expression in zebrafish (measure using *TgBAC(tnfa:GFP)pd1028*) fluorescence 0 days post treatment/infection under labelled conditions. **B.** As A, 1 day post treatment/infection with enlargement showing the increase in the TNF reporter expression in innate immune cells. **C.** As B, 3 days post treatment infection. **D.** Quantification of total fluorescence over time (Mean with 95% confidence interval). Only individual cells exhibiting fluorescence were quantified, therefore excluding central nervous system and gut expression. No statistically significant difference. **E.** Quantification of total TNF reporter expression in innate immune cells at 3 days post infection showing individual infections (Mean with 95% confidence interval). **A-E** 15 treatment/infections per group (5 per 3 times biological repeats). **F.** Change in fungal burden with iNos inhibitor 1400W, and MMF, alone and in combination. DMSO vs. 1400w p=0.047; MMF vs. MMF+1400w p=0.98. **G.** Change in fungal burden with iNos inhibitor LNIL, and MMF, alone and in combination. DMSO vs. LNIL p=0.014; MMF vs. MMF+LNIL p=0.42. **H.** Change in fungal burden with arginase inhibitor Nor-NOHA, and MMF, alone and in combination. DMSO vs. Nor-NOHA p=0.87; MMF vs. MMF+Nor-NOHA p=0.33. **F-H.** Box plots are median, IQR, 5 and 95 whiskers and outliers plotted. Mann-Whitney U test used for significance comparison. Number of infections denoted by n, total from three biological repeats. All infections 500 cfu KN99αGFP.

### Macrophages, but not neutrophil, numbers are reduced after treatment with MMF

In addition to reducing the cytoplasmic pool of BH4, GTP depletion can lead to increased cell death in monocytic cell lines, *in vitro* (Cohn et al., 1999) and we therefore investigated the number of macrophages and neutrophils in MMF treated zebrafish. Macrophage numbers were reduced ∼25% after 24 hours treatment with MMF *in vivo* (Fig. 3A; Fig. S2C; DMSO, median=219; MMF, median=166; P=0.01). We used AZA to distinguish the effect of the GTP depleting properties of MMF and there was no difference in macrophage numbers with AZA treatment, even after 72 hours (Fig. S2A-C). Reduced numbers in macrophages resulted from cell death, as opposed to decreased proliferation, as there were significantly increased numbers of Annexin V positive macrophages observed with MMF treatment (Fig. S2D). In contrast, there was no difference in neutrophil numbers with MMF *in vivo* (Fig. 3B).

**Fig 3.**
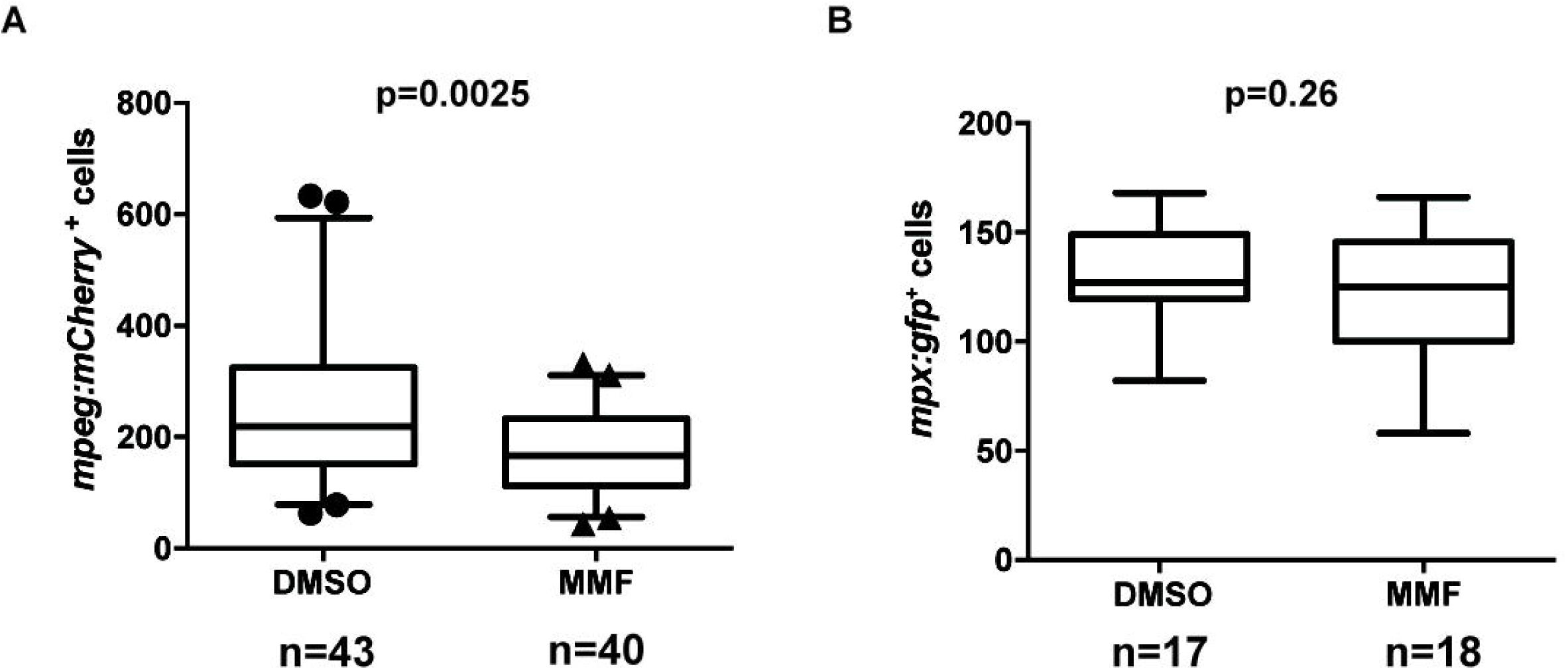
Macrophage, but not neutrophil, numbers were reduced with MMF treatment. A. Number of *mpeg:mCherry* positive cells (macrophages) per larvae counted by flow cytometry. **B.** Number of *mpx:GFP* positive cells (neutrophils) per larvae counted by flow cytometry. Mann-Whitney U test used for significance comparison. Number of larvae analysed denoted by n, total from three biological repeats.

### MMF treatment is associated with reduced phagocytosis early in infection

We have demonstrated previously that the number of macrophages was not the limiting factor in the proportion of crypotoccci phagocytosed by macrophages, and thus early control of cryptococcal infection (Bojarczuk et al., 2016). However, we hypothesised that the reduced macrophage numbers seen with MMF treatment would be sufficient to further limit phagocytosis and increase susceptibility to infection. Using our high content imaging methods (Bojarczuk et al., 2016) we counted the number of macrophages and the number of intracellular and extracellular cryptococci at 1, 2 and 3 days post infection (three biological repeats and a total of 30 infections DMSO and 30 infections MMF). In support of our hypothesis, we found that there was a negative correlation between the number of macrophages and the number of intracellular cryptococci at 24 hours post infection (hpi; Fig. 4B, P<0.0001 R=-0.67). There was no correlation at 24hpi for DMSO treatment alone (Fig. 4A, P=0.06 R=0.00). At 48 and 72hpi there was no correlation between the number of macrophages present and the number of intracellular cryptococci in either DMSO or MMF treated infections (Fig. 4C-F; see legend for P-values). While the number of intracellular cryptococci drives the increase in fungal burden early in infection, due to intracellular proliferation, the majority of growth that contributes to high fungal burden is due to extracellular growth (Bojarczuk et al., 2016). We hypothesised that there would be a consistently higher extracellular burden with MMF treatment due to the restriction of phagocytosis and resulting in the much higher fungal burden seen when there is a transition to extracellular growth. In addition, we predicated that there would not be a correlation between the number of extracellular cryptococci and the total number of cryptococci, as the number of extracellular cryptococci is dependent on the limitation of phagocytosis and, in the case of MMF treatment, the reduced number of macrophages present. Therefore, we compared the proportion of extracellular cryptococci to identify if the high fungal burden seen with MMF later in infection were due to a larger number of extracellular cryptococci early in infection.

**Fig 4.**
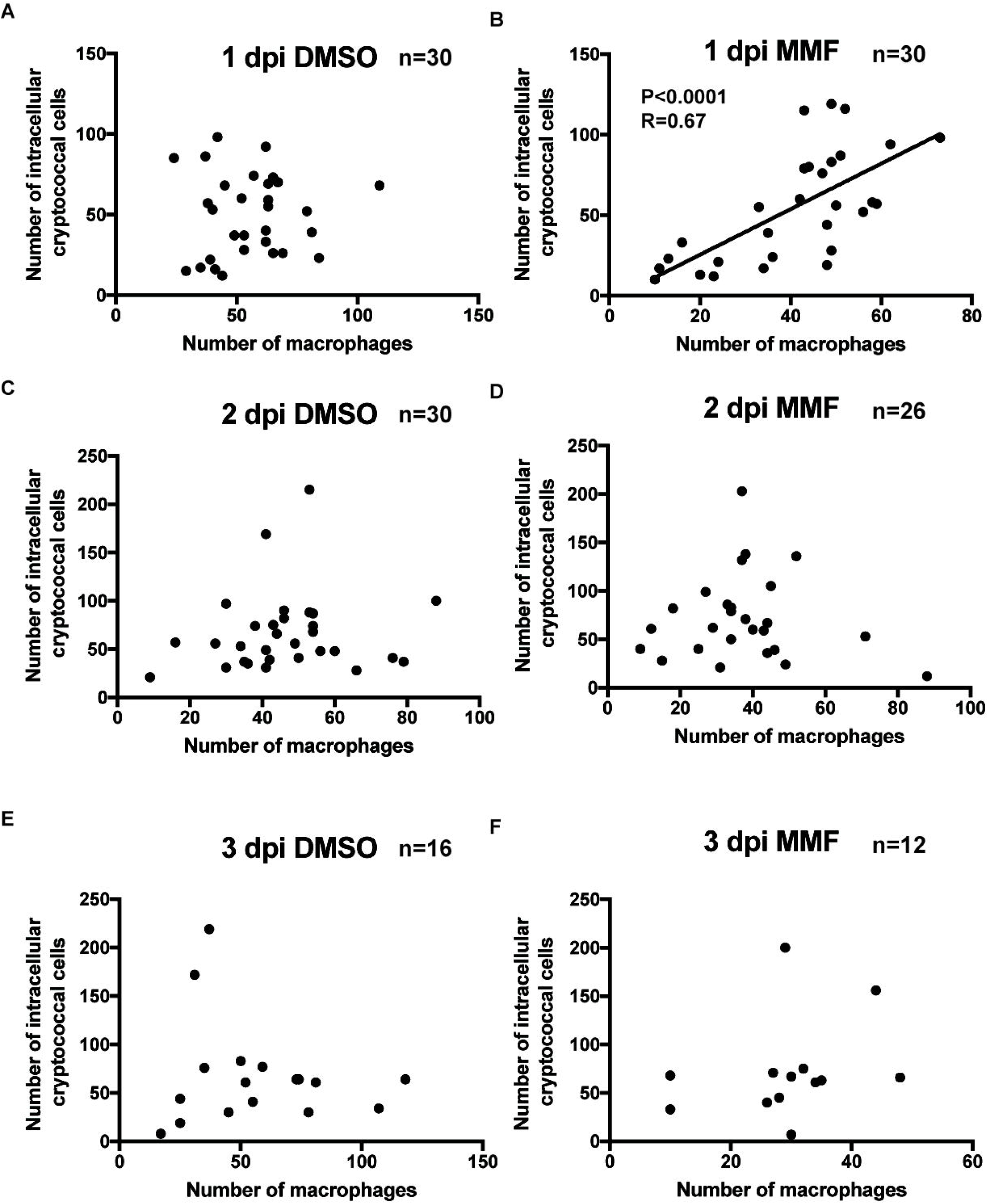
MMF treatment reduced the phagocytosis of *C. neoformans* proportional to the number of macrophages. **A-F.** Linear regression correlation of number of macrophages to the number of intracellular cryptococci. No significant correlation except for B (P<0.0001, R=0.67). Panel title denotes days post infection (dpi) and treatment group. Number of larvae analysed denoted by n, total from three biological repeats. Infections 100 cfu KN99αGFP.

MMF treatment resulted in higher proportion of extracellular cryptococci, with the largest difference at 3dpi, whereas the total numbers of cryptococci were similar at 1 and 2 dpi but diverged at 3dpi (Fig. 5A, B). Linear regression analysis of intracellular or extracellular numbers to the total number of cryptococci gave us a measure of the relative contribution of these populations to fungal burden. We found in all cases that with DMSO treatment there was a linear correlation between the intracellular and total number of cryptococci of greater than R=0.85 (Fig. 5A,C,E). With MMF treatment this correlation was present at 1 and 2 dpi but was lost at 3 dpi, where total numbers of cryptococci were much higher with MMF treatment (Fig. 5B,D,F). However, there was strong correlation with extracellular numbers at 3 dpi with MMF establishment of high fungal burden, demonstrating the switch to extracellular growth with uncontrolled infection (Fig. 6 P<0.0001 R=0.98). Furthermore, this correlation was also present at 2 dpf with MMF, and to a much lesser extent with DMSO treated infections (Fig. 7B, E MMF P<0.0001 R=0.74 DMSO P=0.009 R=0.47). What then was driving the correlation with extracellular numbers at 2 dpi? We analysed the relationship between the proportion of extracellular cryptococci and the total fungal burden. There was no linear correlation in DMSO treated infection (Fig. 6G-I), however MMF treatment again showed a correlation at 2 and 3dpi (Fig. 6K-L). While this was expected at 3 dpi given the dominance of extracellular cryptococci this was not the case at 2 dpi, where the dominance of intracellular growth is still in transition to extracellular growth that ultimately determines the outcome of infection. Taken with the absence of an association with DMSO treatment at 2dpi, we considered what aspects of the macrophage interaction with cryptococci might drive the increase in extracellular cryptococci independent of extracellular growth.

**Fig 5.**
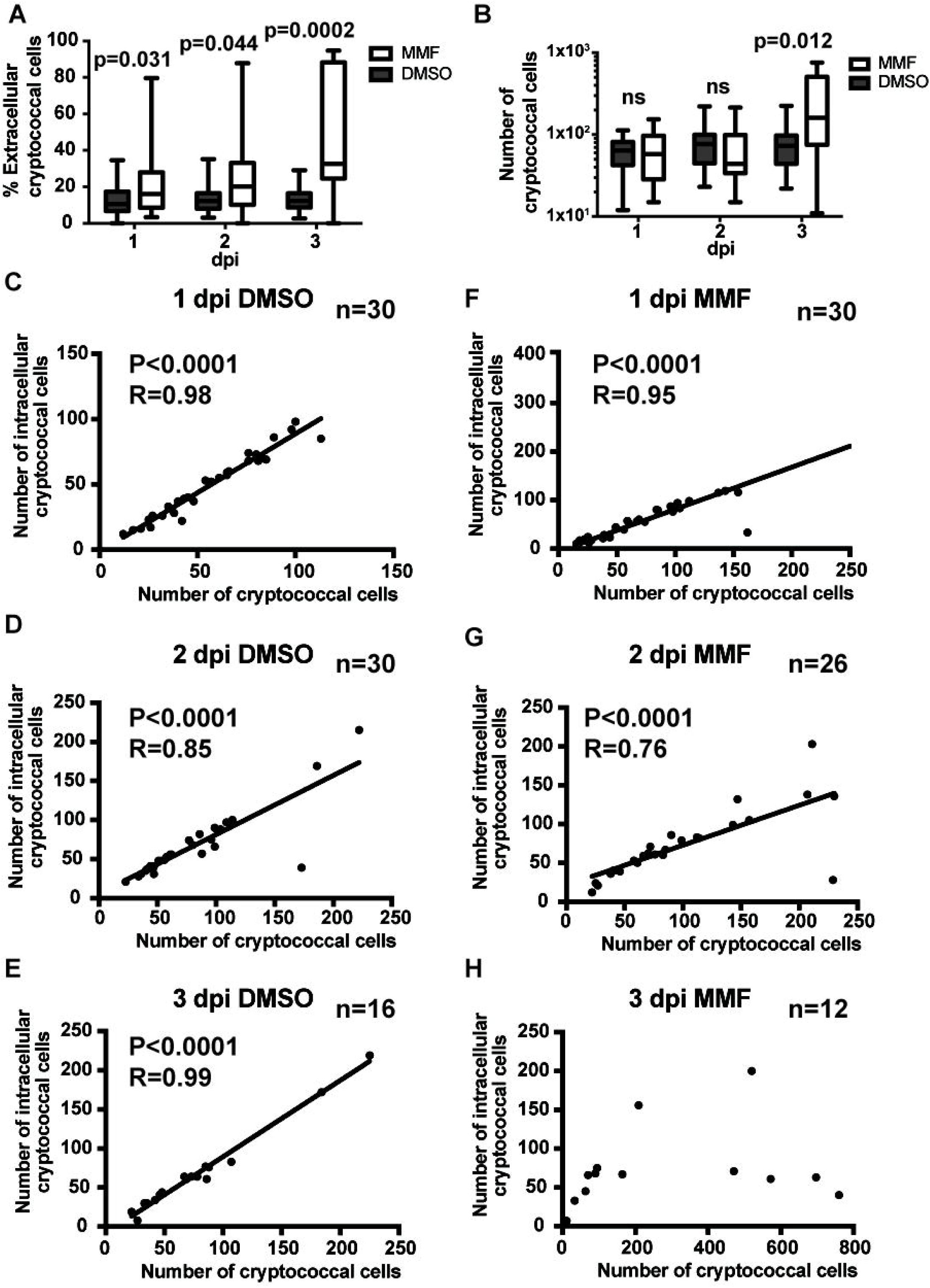
MMF treatment resulted in increased extracellular *C. neoformans* throughout the time course of infection. **A.** The percentage of extracellular cryptococci is increased at all time points with MMF treatment. 30 infections per group (10 per 3 times biological repeats). **B.** The total number of cryptococci is only different at 3 dpi. 30 infections per group (10 per 3 times biological repeats) **C-H.** Linear regression correlation of number of cryptococcal cells to the number of intracellular cryptococci. Panel title denotes days post infection (dpi) and treatment group. Number of larvae analysed denoted by n, total from three biological repeats. Infections 100 cfu KN99αGFP.

**Fig 6.**
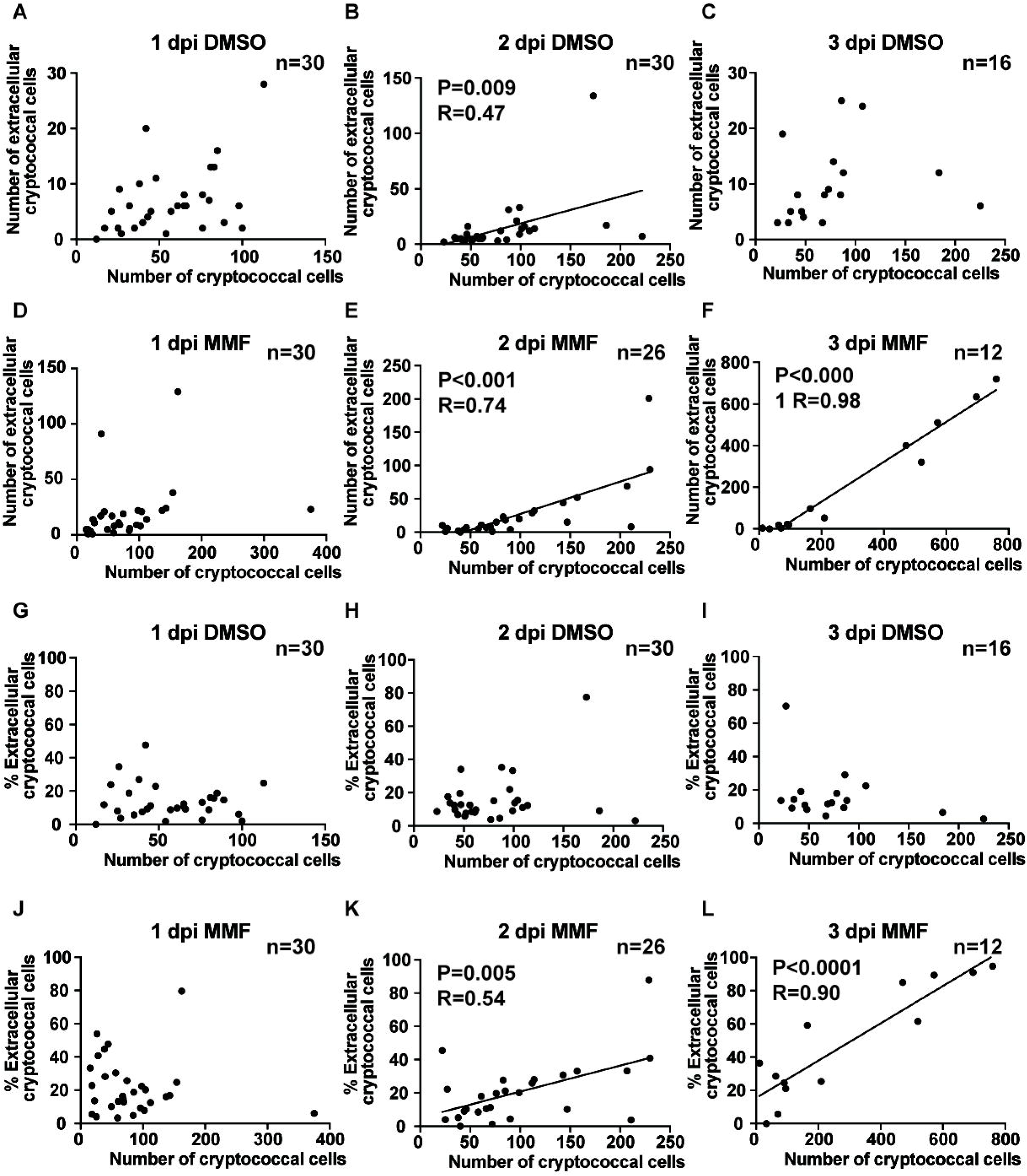
The number and percentage of extracellular *C. neoformans* increasing disproportionately with total fungal burden. **A-F.** Linear regression correlation of number of cryptococcal to the number of extracellular cryptococci. **G-L.** Linear regression correlation of number of cryptococcal to the percentage of extracellular cryptococci. Panel title denotes days post infection (dpi) and treatment group. Number of larvae analysed denoted by n, total from three biological repeats. Infections 100 cfu KN99αGFP.

**Fig 7.**
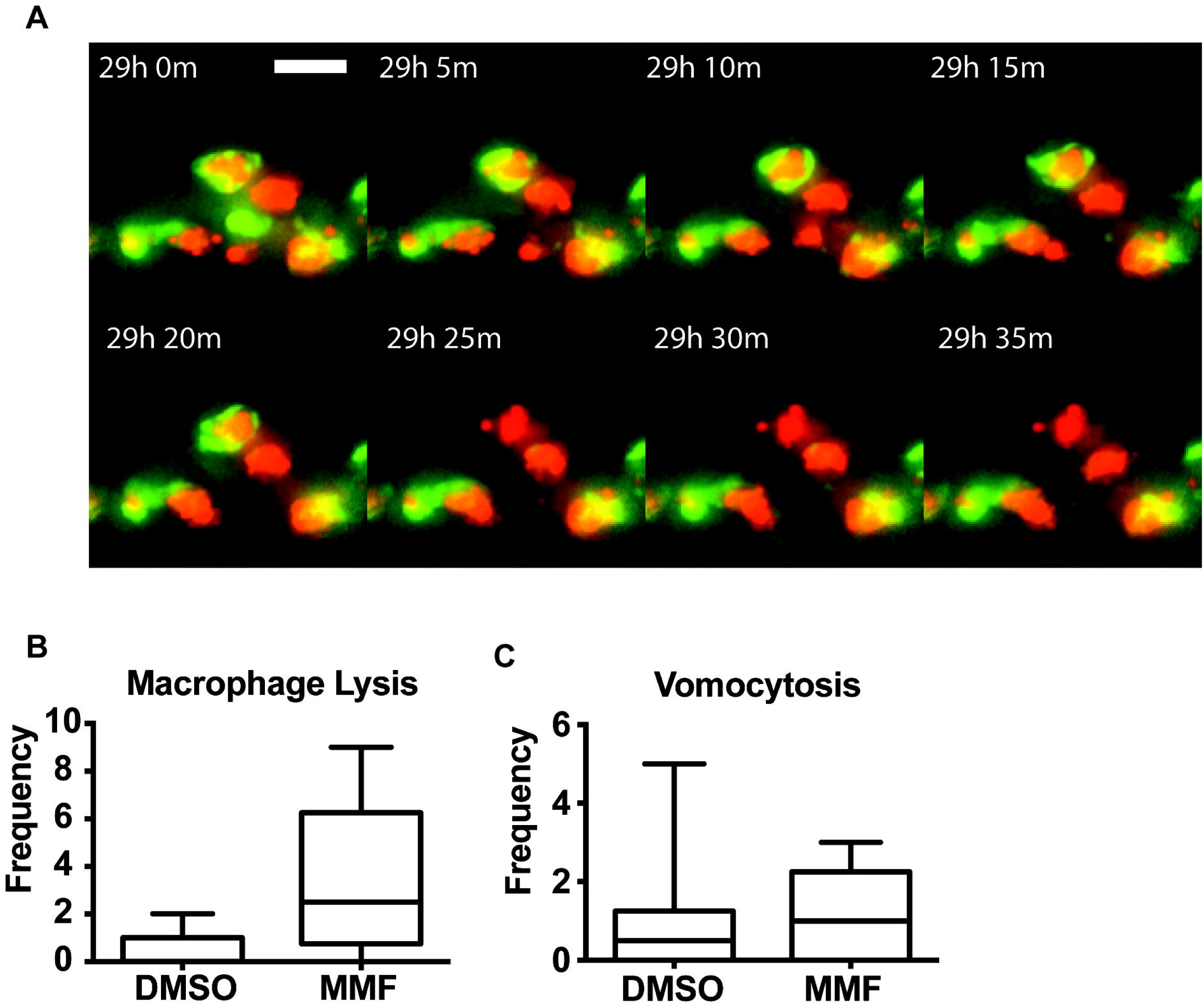
MMF treatment resulted in disproportionate extracellular burden of *C. neoformans* due to macrophage lysis and not vomocytosis. **A.** Example time lapse imaging, from those quantified in B,C, showing macrophage cell lysis and release of intracellular cryptococci at 29 hours post infection. Macrophage (green), (*Tg(mpeg1:Gal4.VP-16)sh256;Tg(UAS:Kaede)s1999t, C. neoformans (red)* KN99αmCherry. **B, C.** Quantification of macrophage lysis and vomocytosis. 12 infections of 500cfu KN99αmCherry per group (4 per 3 biological repeats).

### MMF treatment results in increased release of intracellular cryptococci via macrophage lysis independent of vomocytosis

Cryptococci are known to both lyse macrophages and to escape non-lytically via vomocytosis (Johnston and May, 2013, 2010), both of which would result in the enhanced association between extracellular numbers and fungal burden. Therefore, we used time lapse imaging between 1 and 2 dpi to analyse single cell interactions and measure vomocytosis and macrophage lysis with DMSO and MMF treatment. Using time lapse imaging we identified macrophage cell lysis as the mechanism of increased extracellular cryptococci with MMF treatment. We quantified the number of lysis and vomocytosis events over 12 hours of imaging and found that there was the same number of vomocytosis events with both DMSO and MMF treatment (DMSO, median=0.5; MMF, median=1; P=0.67) but that there was large increase in cell lysis in the MMF treatment group (DMSO, median=0; MMF, median=2.5; P=0.01). Cell lysis was rare with DMSO treatment (lysis in only 3/10 infections analysed) while with MMF treatment lysis occurred in most (8/10 infection analysed) with a mean incidence more than 8-fold per larvae with MMF (MMF, mean=3.5; DMSO, mean=0.4). Thus, the combination of reduced macrophage phagocytosis and release of intracellular cryptococci by macrophage lysis drives the increased fungal burden with MMF treatment.

## Discussion

Here we have shown how the common immunosuppressive mycophenolate mofetil (MMF) causes increased susceptibility to the opportunistic fungal pathogen *Cryptococcus neoformans*. The zebrafish larva is a model of vertebrate innate immunity in the absence of lymphoid cells. Thus, using zebrafish larvae we could show that the effect of MMF occurs in the absence of lymphoid cells, the canonical target of this immunosuppressive. The increased susceptibility to infection is the most serious complication of immunosuppressive therapy and the risks and management of underlying disease must be balanced with the risks of life threatening infection. While calcineurin inhibitors and steroid treatment have been extensively studied, both clinically and using experimental models, the more recently developed anti-proliferative drug MMF is much less understood in the context of infection. Here, we demonstrate that MMF treatment might be considered a larger risk factor for some intracellular pathogens such as *Cryptococcus* and further study is needed to identify potentially important differences in mechanism relating to susceptibility to infection.

Methotrexate is a chemotherapy drug that is used also as an anti-inflammatory. Methotrexate is a competitive inhibitor of dihydrofolate reductase, which leads to depletion of intermediates needed for nucleotide synthesis. In addition to acting as an inhibitor of the folate pathway, the mechanism of methotrexate as an anti-inflammatory drug has been variously proposed to be T cell apoptosis, purine release and cell-cell adhesion. However, methotrexate has recently been shown to act through direct inhibition of the JAK/STAT pathway (Thomas et al., 2015), and this may explain much of its clinical activity in diseases such as Rheumatoid Arthritis. We have shown that MMF treatment results in a specific macrophage phenotype independent of any lymphocyte depletion, hinting that perhaps the effects of MMF might not all be through lymphocyte proliferation as previously thought.

The number of immunosuppressed individuals is increasing, due to both increased life expectancy and adoption of immunosuppression as treatment for an increasing number of diseases (de Haan et al., 2017; Jackaman et al., 2017; Mira et al., 2017; Zeineddine et al., 2016). Macrophages are orchestrators of the immune response, present in almost every organ, with a multitude immune signalling pathways and responses acting through macrophages (Wiegertjes et al., 2016). Therefore, there is a substantial medical need for better understanding immunosuppression mechanisms to identify off target or unintentional suppression of macrophages and to develop new more specific therapies. Here we have demonstrated how *in vivo* models, such as zebrafish, can be used to discriminate different mechanisms of action and identify new aspects of pathogenesis. Furthermore, we and others have demonstrated the use of zebrafish in screening for new therapies for human disease (North et al., 2007; Robertson et al., 2014) and the ability to address both mechanism and drug discovery in the same system makes identification of a new immunosuppressive with greater specificity a realistic aim.

Treatment of cryptococcosis in HIV positive individuals prior to initiation of antiretroviral therapy is now well evidenced, to the extent that cryptococcal antigen testing and prophylactic antifungal treatment is recommended in the HIV positive (Boulware et al., 2014). A major driving force in both cases is the occurrence of immune reconstitution inflammatory syndrome (Longley et al., 2013). Similarly, as stated above, balancing damaging inflammation and progression of infection is a major challenge in the management of those on immunosuppressive therapy. There is large variation in the severity of this problem between individuals (Joseph N. Jarvis et al., 2015). While there is likely to be a significant genetic component to this phenomenon, our data support the suggestion that subclinical or latent infection may be another important factor. Further study is warranted to explore diagnoses and treatment of subclinical or latent infection prior to immunosuppressive treatment, especially where treatment with MMF is otherwise indicated.

We have identified the mechanism of mycophenolate mofetil activity, in contrast to the alternative, drug azathioprine (AZA) due to differences in GTP depletion. AZA produces 6-mecaptopurine (MP) non-enzymatically, which generates the inhibitory metabolites 6-thioguanine (6-TG) and 6-methyl-mecaptopurin ribonucleotides (6-MeMPN). 6-TG inhibits DNA synthesis while MeMPN inhibits *de novo* purine synthesis. Thus, with AZA treatment, cells can still generate GMP, and thus restore GTP levels, through the *de novo* inosine monophosphate (IMP) pathway (depleting cellular ATP). While treatment with AZA will allow the restoration of GMP production, and its downstream products, 6-TG will still have a dominant effect on DNA synthesis. In contrast, MMF inhibits inosine monophosphate dehydrogenase enzyme (IMPDH) required for the *de novo* IMP pathway. Therefore, with MMF treatment GMP can only be generated through the salvage pathway increasing the competition for GMP (mainly from nucleotide synthesis) results in depletion of the overall GTP pool. The GTPases, G protein coupled receptors (GPCRs), and the small and large GTPase families all rely on availability of GTP for activation, suggesting possible mechanisms by which this might regulate macrophage function. Chemokine receptors are GPCRs with critical roles in regulating macrophage function (Bhatia et al., 2012). The Rho family of small GTPases are essential for the regulation of the actin cytoskeleton (Niedergang and Chavrier, 2005). Inhibition of the Rho family GTPase Rac1 promotes apoptosis of monocyte derived osteoclasts (Fukuda et al., 2005) but Rac1 activation is required for *Salmonella enterica* var. Typhimurium induced apoptosis of macrophages (Forsberg et al., 2003). In addition, it is likely that depletion of GTP will result in vesicle trafficking defects as many of its steps are regulated by small GTPases (Niedergang and Chavrier, 2005). Finally, GTP is a key metabolite linking the citric acid cycle to gluconeogenesis, and is necessary for the glycosylation of proteins.

The inflammatory, or classical, activation of macrophages is the best characterised and evidenced route for the clearance of cryptococcal infection (Joseph N Jarvis et al., 2015; Leopold Wager et al., 2014). The iNOS pathway is critical in classical activation of macrophages through the direct production of antimicrobial reactive nitrogen species and the positive feedback signalling loop of pro-inflammatory cytokines gamma interferon and tumour necrosis factor alpha (TNFalpha; (Jacobs and Ignarro, 2003; Sandau et al., 2001)). Therefore, we hypothesised that reduced cellular GTP would alter macrophage derived TNFα levels and any iNOS mediated control of cryptococcal infection. While we found that *C. neoformans* induced TNFα *in vivo* in zebrafish, MMF had no effect on this response. Disruption of TNFα signalling in murine models increases susceptibility to infection (Hage et al., 2003). *C. neoformans* induced TNFα expression is measurable as early as two days post infection in the murine lung but using a fluorescent protein transcription reporter we were able to measure expression at 24 hours post infection (Herring et al., 2002; Huffnagle et al., 1996). TNFα is required for protective adaptive immunity (Huffnagle et al., 1996) but is not sufficient for protective macrophage responses to cryptococcal infection and, with the absence of TNFα modulation, we concluded that restriction of TNFα is not a factor in increased susceptibility of the innate immune system to cryptococcal infection with MMF. GTP is a precursor for the iNOS enzymatic cofactor tetrahydrobiopterin (BH4) and therefore GTP depletion can limit the activity of the iNOS pathway. While the requirement for iNOS/classical activation of macrophages has been well studied in the context of T cell activation of macrophages we have addressed the question of iNOS activity in the absence of T cells *in vivo*. We found that pharmacological inhibition of iNOS had a small effect on susceptibility to cryptococcal infection, with a significant but slight increase in fungal burden. Arginase is an enzyme that competes for the iNOS substrate L-arginine and inhibition of arginase will enhance iNOS activity. However, we found that arginase inhibition had no effect on cryptococcal fungal burden supporting our conclusion that there was little activation of the iNOS pathway with cryptococcal infection in the absence of adaptive immunity.

We identified that macrophage numbers were reduced with MMF treatment and we identified increases in macrophage cell death both with and without infection *in vivo*. MMF has been described as reducing infiltration of macrophages in rodent models (Jiang et al., 2012; von Vietinghoff et al., 2011). However, these studies could not differentiate reduced recruitment of macrophages from macrophage cell death and our data suggest that, while there may be reduced macrophage recruitment, macrophage cell death is a significant factor in any reduction in macrophage numbers with MMF treatment. A study of the anti-inflammatory properties of MMF in lung reperfusion injury in a rat model showed reduced brochoalveolar lavage cell counts and reduced macrophage chemotactic protein 1 (MCP-1/CCL2). As MCP1 is primarily a macrophage derived chemokine this supports our data that MMF may have a direct effect on tissue macrophage numbers. Clinical monitoring of tissue macrophage number and function is very difficult but we considered that differential cell counts from bronchoalveolar lavage of patients on MMF versus AZA represented one of the very few cohorts that might be studied. However, we could only identify a single existing analysis of this kind (Speich et al., 2010) and concluded that, as such patients will have widely variable clinical presentation and treatment history, controlling for the different confounding factors would require a very large sample size and a retrospective meta-analysis of a large number of smaller studies would be additionally hampered by differences in reporting and data collection.

High dose mycophenolate has been demonstrated to have antimicrobial action against *C. neoformans* and GTP depletion in fungal cells was suggested as a potential drug target (Morrow et al., 2012). However, our work suggests that clinical use of MMF for this indication is would be highly inadvisable due to increased susceptibility to cryptococcosis and, likely, other intracellular opportunistic pathogens. The dose of MMF used in our study is below the known patient serum concentration range for mycophenolic acid (0.5μM versus 3-11μM). In contrast, the lowest dose of mycophenolic acid with antimicrobial activity tested was 16μM and was only used *in vivo* in a nematode model where there is no cell mediated immunity. Given the clear similarity between fungal and mammalian inosine monophosphate dehydrogenase active site (Morrow et al., 2012) there appears to be little possibility that there might be a fungal specific inhibitor, unlike for the fungal calcineurin, which is a viable drug target (Juvvadi et al., 2017).

In contrast to cryptococcal fungal burden, we found no effect on mycobacterial burden. Macrophage depletion can increase susceptibility to mycobacterial infection in zebrafish with clodronate liposomes and in the csf1r mutant (Pagán et al., 2015). Our experimental model of MMF treatment only has comparable macrophage depletion to the lowest dose of clodronate liposomes at two days post treatment. Liposomal clodronate rapidly depletes macrophages and we have shown previously that there is greater than 50% depletion six hours post treatment and this agrees with mammalian studies where depletion occurs within hours (Bojarczuk et al., 2016; Camilleri et al., 1995). Similarly, the csfr1 mutant has supressed macrophage numbers from the initiation of infection. In the absence of adaptive immunity the innate immune system in zebrafish will efficiently phagocytose mycobacteria (unlike cryptococci) and there is recruitment of uninfected neutrophils and macrophages to infected cells. The continued recruitment of uninfected macrophages is required for control of mycobacterial infection in zebrafish as depletion of uninfected macrophages (by clodronate or in the csfr1 mutant zebrafish) increases susceptibility to infection. A proposed mechanism for this requirement is the phagocytosis of extracellular mycobacteria released following infected macrophage or neutrophil cell death (Hosseini et al., 2016). This is distinct from the progression of cryptococcal infection where there is very little macrophage lysis (discussed below) and where phagocytosis of extracellular fungal cells in limited by their polysaccharide capsule, not by the number macrophages (Bojarczuk et al., 2016). Therefore, there we did not find an increase in mycobacterial burden with MMF treatment because there was insufficient rate in macrophage depletion compared to liposomal clodronate treatment (or the existing defect in csfr1 mutant zebrafish) and that any increase in macrophage lysis was not significant when compared to the rate of infected macrophage cell death normally seen in mycobacterial infection of zebrafish.

We have previously demonstrated that macrophage numbers are not the limiting step in the control of cryptococcal infection. However, here we show that as macrophages are depleted there is a limitation in phagocytosis independent of the cryptococcal population. The initiation of the immune response to infection relies on tissue resident macrophage populations. While neutrophils will rapidly chemotax from the circulation to wound sites, macrophages are resident in tissue for pathogens that enter through e.g. the respiratory system, or avoid and/or are missed the immune response at a wound. A tight threshold on immune cell depletion leading to disease is well known, most notably for CD4 cell counts in those with AIDS. Finally, we identified a large increase in the release of intracellular pathogens following MMF treatment via macrophage lysis. Macrophages are known to use apoptosis to kill intracellular pathogens (Dockrell et al., 2003). However, there is no evidence that macrophages use apoptosis in the control of cryptococcal infection. We found that the release of intracellular cryptococci was significant driver of the increase in fungal burden seen with MMF treatment.

Thus, we have shown that, independent of lymphoid cells, mycophenolate mofetil causes macrophage depletion and lysis, increasing susceptibility to opportunistic fungal infection. This highlights the importance of understanding the mechanism of drug action, especially critical for those on immunosuppressive therapy where there is a small therapeutic window for benefit versus risk of infection, and that for many immunosuppressive drugs in clinical use, further research is needed to inform the relative benefits and risks of treatment.

## Acknowledgements

We thank Arturo Cassadevall for the gift of 18B7 antibody, Peter Williamson for the kind gift of mCherry amplified from pKUTAP and the Bateson Centre aquaria staff for their assistance with zebrafish husbandry. SAJ, RHG, RJE, RH, AB and AL were supported by Medical Research Council and Department for International Development Career Development Award Fellowship MR/J009156/1. SAJ was additionally supported by a Krebs Institute Fellowship, Medical Research Foundation grant R/140419 and Medical Research Council Centre grant (G0700091). RHG was supported by a Bateson Centre Biomedical Science scholarship. RH was supported by a Colin Beattie Biomedical Science scholarship. RJE was supported by a British Infection Association Fellowship. PME was supported by a Wellcome Trust and Royal Society Sir Henry Dale fellowship. SAR was supported by a Medical Research Council Programme Grant (MR/M004864/1). RCM was supported by a Lister Fellowship. RCM and EB were supported by the European Research Council Consolidator Award “MitoFun”.

---

**Fig. S1. Treatment with Mycophenolate mofetil does not affect mycobacterial burden. A, B.** Representative fluorescence image of the median mycobacterial burden for DMSO versus MMF. **C.** Quantification of bacterial burden. 84 infections from 2 biological repeats.

**Fig. S2. Macrophage depletion does not occur with azathioprine. A.** Number of *mpeg:mCherry* positive cells (macrophages) per larvae counted by flow cytometry at 3 days post treatment with MMF. **B.** Number of *mpeg:mCherry* positive cells (macrophages) per larvae counted by flow cytometry at 3 days post treatment with AZA. **C.** Time course of changes in Number of *mpeg:mCherry* positive cells (macrophages) per larvae with MMF treatment. **D.** Quantification of the number of Annexin V positive macrophages. Mann-Whitney U test used for significance comparison between macrophage counts and Fisher’s exact test used to compare proportion of macrophages Annexin V positive. Number of larvae analysed denoted by n, total from three biological repeats.

**Fig. S3. Generation and charctrisation of KN99αGFP and KN99αmCherry. A.** To obtain fluorescently labeled *C. neoformans* serotype A, strain KN99 α was biolisticly transformed (Toffaletti et al., 1993; Voelz et al., 2010) with a plasmid pAG32_GFP (Voelz et al., 2010) encoding for a green fluorescent protein (GFP) or with a plasmid pRS426H-CnmCherry encoding for a *C. neoformans* - codon optimized red fluorescent protein mCherry. Diagram representing the plasmid p RS426H. Abbreviations: H3p, Histone 3 promoter; H3t, Histone 3 terminator; Act1p, actin promoter; HygR, Hygromycin resistance gene; Trp1t, trp1 terminator. Figure was made using Dog 2.0 software (Ren et al., 2009). **B. C.** Stable transformants were tested for their sensitivity against several stress conditions mimicking hostile environment inside the phagocytes. Three independent experiments were carried out where the transformants grew up at 37°C and 5% CO_2_ without shaking in the presence/absence of H_2_O_2_ or NaCl mimicking oxidative or cell wall stresses, respectively (Voelz et al., 2010). After 24 hours of growth serial dilutions of surviving cells were plated onto YPD plates and colony-forming units (CFU) were counted after 0 and 48 hours growth at 25°C incubator. CFUs relative to time point 0 were calculated. **D.** GFP and mCherry expressing transformants were tested towards their survival and intracellular proliferation rates (IPR) inside J774 macrophages. For IPR 1 ml of the J774 macrophage-like cell lines (0.5 x 10^5^ cells/ml) were infected at MOI 1:10 with a wild type KN99α and GFP or mCherry-expressing transformants (0.5 x 10^6^ cells/ml) opsonized with 18B7 antibody (a kind gift of Prof. Arturo Casadevall, Johns Hopkins Bloomberg, School of Public Health, Baltimore, United States of America) as described previously (Ma et al., 2009; Voelz et al., 2010). The IPRs were assessed after initial 2 and following 24 hours of infection. P-values of statistical analysis (Wilcoxon matched-pairs signed rank test) of results from CFU counts from stress treatments of GFP/mCherry-positive strains compared to a parental strain KN99.

